# Protein Translation Can Fluidize Bacterial Cytoplasm

**DOI:** 10.1101/2024.01.16.575857

**Authors:** Palash Bera, Abdul Wasim, Somenath Bakshi, Jagannath Mondal

**Affiliations:** Tata Institute of Fundamental Research, Hyderabad 500046, India; Department of Engineering, University of Cambridge, Cambridge, United Kingdom

## Abstract

The cytoplasm of bacterial cells is densely packed with highly polydisperse macromolecules that exhibit glassy dynamics. Research has revealed that metabolic activities in living cells can counteract the glassy nature of these macromolecules, allowing the cell to maintain critical fluidity for its growth and function. While it has been proposed that the crowded cytoplasm is responsible for this glassy behavior, a detailed explanation for how cellular activity induces fluidization remains elusive. In this study, we introduce and validate a novel hypothesis through computer simulations: protein synthesis in living cells contributes to the metabolism-dependent fluidization of the cytoplasm. The main protein synthesis machinery, ribosomes, frequently shift between fast and slow diffusive states. These states correspond to the independent movement of ribosomal subunits and the actively translating ribosome chains called polysomes, respectively. Our simulations demonstrate that the frequent transitions of the numerous ribosomes, which constitute a significant portion of the cell proteome, greatly enhance the mobility of other macromolecules within the bacterial cytoplasm. Considering that ribosomal protein synthesis is the largest consumer of ATP in growing bacterial cells, the translation process likely serves as the primary mechanism for fluidizing the cytoplasm in metabolically active cells.

## INTRODUCTION

In the absence of cytoskeletal filaments and motor proteins, biomolecular reactions within the bacterial cytoplasm are predominantly constrained by diffusion. Recent experiments involving single molecule tracking in living cells have highlighted the prominent role of diffusion-limited search times as the determining factor for reaction kinetics [1–4]. As a result, comprehending the mechanisms that influence the diffusional properties of various cytoplasmic components is crucial for grasping the spatiotemporal organization of biomolecular reactions within individual bacterial cells.

The dynamics of various cellular entities in the bacterial cytoplasm exhibit significant variability and frequently deviate from simple diffusion. The bacterial cytoplasm represents a densely populated environment comprising a multitude of polydisperse entities [5–10], ranging in size from a few nanometers (such as small proteins, ribosomes, metabolites, etc.) to micrometers (including DNA, polysomes, large proteins, etc.). This polydisperse crowding has been demonstrated to influence the dynamics of individual components in a size-dependent manner [11]. A recent study by Parry et al. [12] further reveals that the mode of dynamics also relies on the size of the particle under examination. In aggregate, their findings suggest that the bacterial cytoplasm displays dynamics reminiscent of glass-like behavior, where smaller components exhibit liquid-like dynamics and larger components exhibit solid-like dynamics.

The highly crowded and polydisperse nature of the cytoplasm is expected to underlie the size-dependent glassy dynamics observed in cytoplasmic particles. At significantly high crowder densities, particles can become entrapped and unresponsive to external perturbations, resembling solids. While in a monodisperse system, this scenario could lead to crystallization, in a complex and polydisperse system like the bacterial cytoplasm, the dynamics turn chaotic and glassy, which aligns with observations made by Parry et al [12].

However, comprehending the mechanistic intricacies governing these glassy dynamics would benefit from computational investigations, which enable precise control and consideration of confounding perturbations. This contrasts with experimental settings where apparently straightforward perturbations might trigger various unpredictable changes in biomolecular states, complicating the identification of intended direct effects. In this study, we endeavor to address this challenge using a Brownian dynamics model of the bacterial cytoplasm. This model allows for systematic exploration of diverse factors that could impact the dynamics of cytoplasmic components.

In the paper by Parry et al. [12], they also discovered that in metabolically active cells, the dynamics of larger components are fluidized by cell metabolism, potentially playing a critical role in determining their enzymatic activity. However, the mechanism underlying the metabolic activity-dependent fluidization of the glassy cytoplasm remains unclear. An earlier investigation by Weber et al. [13] demonstrated that ATP-dependent non-equilibrium fluctuations contribute to the macro-molecular motion in cells, enhancing their mobility. In line with this observation, Parry et al. identified ATP as a pivotal factor in fluidizing the dynamics of various cytoplasmic components. Numerous potential mechanisms of ATP-dependent fluidization have been proposed (including protein conformational changes, non-equilibrium fluctuations, fluid displacement, etc.). However, to align with experimental observations, such a mechanism should exhibit strong ATP dependence and induce fluidization or glassification of cytoplasmic components in a size-dependent manner.

In this study, we propose a hypothesis that underscores the role of ATP-dependent translation activity in metabolically active cells as the primary force for cytoplasmic fluidization. Translation stands as the primary ATP consumer within the cell, with ribosomes being the central players which are large molecular machines highly concentrated in the cytoplasm [10, 14]. The presence or absence of ATP could shift the state of these abundant molecular machines from primarily engaged in translation to predominantly free diffusion [8, 10]. Through computer simulations, we delve into how macromolecular crowding triggers glassy dynamics in the cytoplasm and assess whether the dynamic fluctuations in ribosome diffusivity, dictated by their activity, could sufficiently perturb the remaining cytoplasm to induce fluid-like motion.

Our simulations demonstrate that molecular crowding indeed leads to size-dependent glassy dynamics within the cytoplasm. Subsequently, we develop a model to describe ribosome state switching during translation (initiation: fast to slow and termination: slow to fast). Our findings highlight that the substantial changes in the dynamics of numerous ribosomes during translation initiation and termination can effectively disrupt the motion of other cytoplasmic entities, promoting fluidic behavior. To provide further substantiation, we conduct additional computational and bioinformatics investigations to elucidate why ribosomes and their biochemical states are uniquely suited to serve as the primary driving force behind the transformation of the inherently glassy cytoplasm.

## RESULTS AND DISCUSSION

### A. Modelling the effects of molecular crowding in the bacterial cytoplasm

We initiate our exploration by creating a model of the bacterial cytoplasm to investigate the effects of various configurations on molecular dynamics. The bacterial chromosome is represented as a ring polymer with interconnections between segments, established using inter-gene contact probabilities derived from Hi-C experiments [15]. This is combined with a polymerphysics-based framework within a spherocylindrical confinement (see the Model and method section for further details).

To simulate an excessively crowded condition resembling the bacterial cytoplasm, the simulated cell is densely packed with monomeric ribosomes (50S and 30S) as well as 13-bead polymer chains representing 70S ribosomes, referred to as polysomes. This is set in an 80:20 polysome-ribosome ratio [8, 10]. While ensuring bonded connectivity within the chromosome, we employ soft excluded volume interactions to model the mutual interactions among all species coexisting within the cell. The dynamics of each constituent particle within this densely populated cell adhere to Brownian dynamics. All length and time measurements are described in reduced units relative to the diameter of the DNA bead (*σ*) and Brownian time (*τ*_*B*_), respectively (as detailed in the Model and methods section).

In Figure 1(a), a representative snapshot of *E. Coli* cy-toplasm is presented, with color-coded beads representing various particles. Additionally, for enhanced visualization of their spatial distribution, we have individually plotted each of the components (Figures 1 (b), (c), (d) and (e)).

**FIG. 1.**
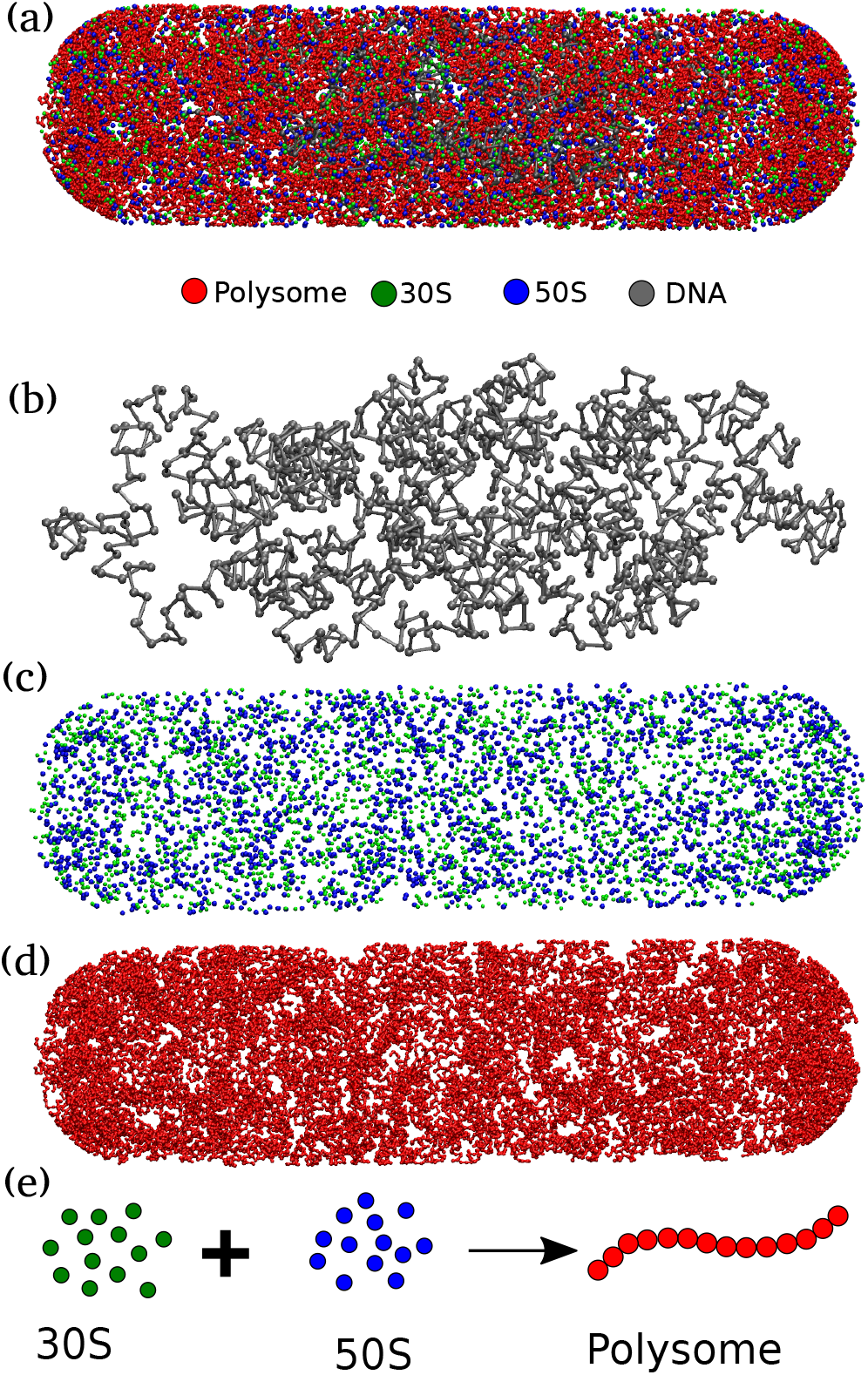
A representative snapshot of the model bacterial cytoplasm. The upper figure (a) depicts a snapshot of the bacterial cytoplasm, enclosed by the cell wall, where different types of particles are distinguished by color-coded beads. To enhance visualization, we extracted the distinct particle types from the snapshot and presented them individually (from top to bottom: DNA, 30S and 50S combined, and polysomes). The lower schematic illustrates two states of ribosomes: 1. separated into 50S and 30S monomers, and 2. combined into 70S and integrated within a polysome.

As affirmed by superresolution experiments [8–10] and validated by our previous computational model [8, 16], our current model consistently illustrates a significant spatial separation between ribosomes and DNA, as shown in Figures 2 (a-b). In an effort to preserve configurational entropy, DNA tends to condense away from the cell walls, consequently creating distance between itself and large particles like ribosome subunits and ribosome polymers. These ribosome particles are predominantly situated within the end-cap region of the bacterial cytoplasm, an area where DNA is relatively sparse. This segregation phenomenon is notably pronounced for ribosome chains, known as polysomes.

**FIG. 2.**
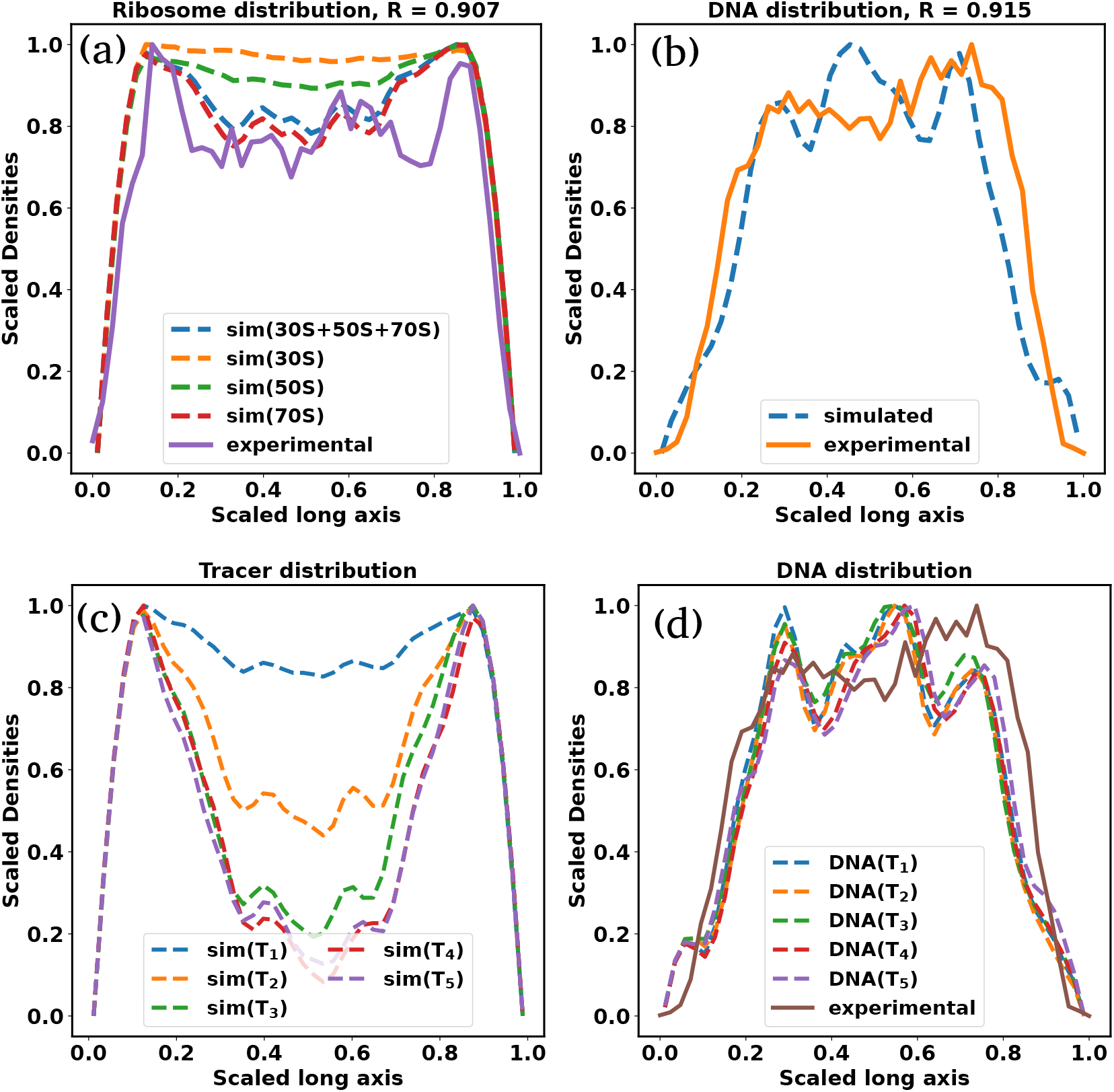
Size dependent spatial distribution of cytoplasmic particles. The figure shows the linear density profiles of ribosomal subunits (a) and DNA (b). Here, solid lines correspond to experimental data, while dashed lines represent simulated densities. For both DNA and ribosomal subunits, a high Pearson correlation coefficient (R) between the experimental and simulated densities indicates strong agreement between our simulations and experimental observations. Notably, the figure clearly demonstrates significant spatial separation between ribosomes and DNA. Figures (c) and (d) depict the linear density profiles of tracer particles (*T*_1_ *− T*_5_) and DNA. Similar to ribosomes, large tracer particles also exhibit noticeable segregation from the nucleoid, with the extent of this segregation being contingent on their respective sizes. However, the presence of tracer particles does not lead to additional compaction of the nucleoid.

When comparing the spatial distribution profiles of polysomes to the nucleoid, a clear segregation pattern emerges. This pattern closely mirrors experimental results obtained under relevant growth conditions, as documented in Mohapatra’s studies [9]. This finding aligns with our earlier work’s polysome-nucleoid demixing hypothesis [8, 16]. Notably, polysomes, being chains of ribosomes, exhibit a lower likelihood of intermixing with DNA chains when compared to individual 30S and 50S ribosome subunits. This reduced intermingling is due to the significant loss of configurational entropy involved.

### B. Size dependent spatial distribution of cytoplasmic particles

In the following investigation, we delve into the spatial distribution of cytoplasmic particles in relation to the nucleoid and ribosomes within our model cytoplasm. Our aim is to ascertain whether the distribution of these particles correlates with their individual sizes. To explore this, we introduced five distinct tracer particles, each characterized by varying diameters (*T*_1_ = 10 nm, *T*_2_ = 20 nm, *T*_3_ = 30 nm, *T*_4_ = 40 nm, and *T*_5_ = 50 nm). These cytoplasmic particles interact with each other and the rest of the cytoplasmic components through excluded volume interactions (see Method and SI).

In Figure 2(c) we present a comparison of the individual particle distributions along the long axis of the bacterial cytoplasm. The density profile of the nucleoid, as displayed in Figure 2(a), serves as a reference point for assessing the relative segregation of these tracer proteins from the nucleoid. Much like ribosomes, large tracer particles also exhibit noticeable segregation from the nucleoid, and the extent of this segregation depends on their respective sizes.

Importantly, the presence of tracer particles does not lead to additional compaction of the nucleoid (Figure 2(d)). However, their spatial arrangement is indeed influenced by the nucleoid’s presence. The compacted nucleoid seems to act as a sieve, effectively segregating the proteins from its core. Consequently, proteins are predominantly located in the end-cap region of the cytoplasm. Notably, larger tracer particles are strongly excluded from the nucleoid and tend to congregate in the ribosome-rich regions at the cytoplasm’s end cap.

Considering that the experimental research has demonstrated that only larger tracer particles typically exhibit glassy dynamics, our subsequent analysis zeros in on particles exceeding 50 nm in size. As evidenced in Figure 2(c), these larger particles are exclusively concentrated within the ribosome-rich regions of the end cap. Consequently, our primary focus lies in the dynamic analysis of particles situated within the end-cap region. Accordingly, the following sections consider the analysis of particle dynamics in a cubic volume representative of the composition and configuration of the ribosome-rich end cap of the cytoplasm.

### C. Molecular crowding causes size-dependent glassy dynamics of proteins in cytoplasm

To scrutinize the impact of molecular crowding on observed dynamics, we conduct a comparative analysis of the dynamics of individual components in the presence and absence of all other cellular constituents. To comprehensively account for the diverse range of crowding species found in the bacterial cytoplasm, it is imperative to consider additional crowding particles apart from the nucleoid and ribosomes. However, attempting to simulate the entire bacterial cytoplasm with the vast number of crowding entities poses significant computational challenges, rendering simulations excessively sluggish and unwieldy.

Therefore, given our earlier findings that larger particles are predominantly situated in the end-cap region, our present focus centers on detailed simulations of a representative volume sharing similar dimensions and composition with the cytoplasmic end cap. In order to achieve a high packing fraction representative of the cytoplasm (pf = 0.57), we introduce five distinct types of polydisperse protein particles (*P*_1_ *− P*_5_) with sizes ranging from 13 to 18 nm. Moreover, we have also simulated them individually. For these simulations, we have chosen a very low copy number (100) of each individual monomeric component to ensure that the components dont become overcrowded by themselves, while still providing sufficient data for statistical analysis of dynamic properties. We compare the dynamics of individual components in the presence and absence of all the other components. Detailed descriptions of the models and corresponding calculations of mean-squared displacements can be found in the Model and methods section.

In the absence of any crowding agents, the dynamics of the proteins and tracer particles exhibit super-diffusion on a remarkably short timescale, while also demonstrating the expected linear scaling of MSD at a very long time scale (Figure 3(b)). When these moieties are placed in the crowded environment of the simulated cytoplasm, their dynamics can no longer be described as simple diffusion. Even for the smallest proteins, the mean squared displacement slows down over longer periods of time. For proteins and tracer particles (*T*_5_) a clear presence of an intermediate plateau is evidenced and indicates the glass transition (Figure 3 (c) and (d)). At shorter timescales, these moieties explore the local environments in a superdiffusive manner without being aware of the crowded neighborhood. At the intermediate timescales, they encounter immediate neighborhood obstacles and become caged. At longer timescales they can escape the cage and their motion mimics an effective diffusive motion, which is much slower than their thermal diffusion constant in this media. This two-step dynamic is a hallmark of glass transition and is seen in the entire range of particles explored here.

**FIG. 3.**
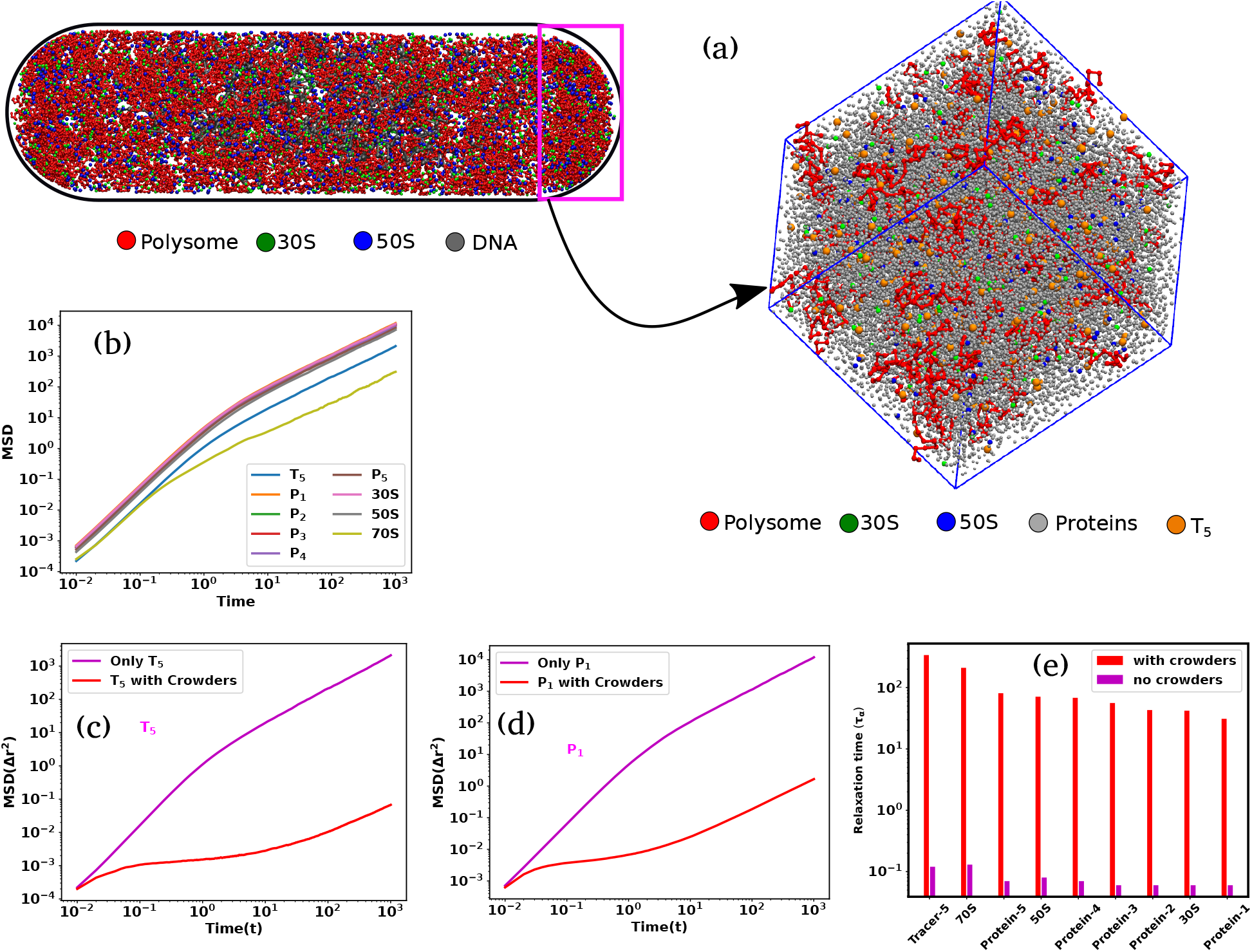
Size-dependent glassy dynamics of particles in a crowded cytoplasm. (a) The end cap of the cells is highlighted by the small rectangle. Considering a cubical box with a volume akin to that of the cell’s end cap, we have inserted the different types of cytoplasmic particles inside it. (b) MSD as a function of time for different components in our simulations in the absence of crowders. (c-d) The MSD for different particle sizes is compared in the presence and absence of crowders for *T*_5_, and *P*_1_, respectively. In the presence of crowders, all the components show a ballistic region and diffusive region in very short and longer time scales respectively and these two regions are separated by a plateau which is an indicator of glassy dynamics. (e) Bar plots of structural relaxation time for different sizes of particles both in the presence and in the absence of the crowders. The relaxation times are larger in the presence of crowders, and this effect is also dependent on particle size.

To further investigate the potential relaxation dynamics typically associated with a glassy matrix, we calculate the structural relaxation time (*τ*_*α*_) for particles of varying sizes, both in the presence and absence of crowding agents. This calculation involves assessing the overlap function (for detailed information, refer to the Material and Methods sections). The relaxation time is typically determined from a two-point density-density autocorrelation function. This parameter provides valuable insights into how the system gradually loses its memory over time. In Figure 3(e), we present bar plots illustrating (*τ*_*α*_) for particles of different sizes in ascending order, denoted by red (in the presence of crowding) and magenta (in the absence of crowding) colors. Notably, in the presence of crowding, the relaxation times for all particles are significantly extended compared to their values in the absence of crowding. Furthermore, the relaxation times are notably influenced by the particle sizes, particularly when crowding is present. These observations strongly suggest that the glassy dynamics, when crowding is a factor, become more pronounced for larger particles.

### D. Dynamic heterogeneity and non-Gaussian distribution of displacements

Beyond just looking at the overall behavior of the tracer particles as a group, as evidenced from mean squared displacements, examining the individual paths of these particles clearly reveals that their dynamics is specifically influenced by the molecular crowding in the system. As an illustration, we have plotted the trajectories of the (*T*_5_) particles in (*yz*) plane in the absence (Figure 4 (a)) and in the presence (Figure 4 (c)) of crowders respectively. The start and end point of these trajectories are marked by magenta and green circles. In the absence of any crowder, the shape of individual trajectories are reminiscent of Brownian motion and sample a large space uniformly. However, in the presence of crowders, the variation of trajectory sizes of *T*_5_ within the same time period shows that there are co-existence of fast (blue) and slow (red) moving particles. Such behaviour is a signature of dynamic heterogeneity (DH). DH is a prominent feature of glassy dynamics, indicating significant variations in local particle dynamics. It implies the coexistence of regions with both highly mobile and less mobile particles [17, 18].

**FIG. 4.**
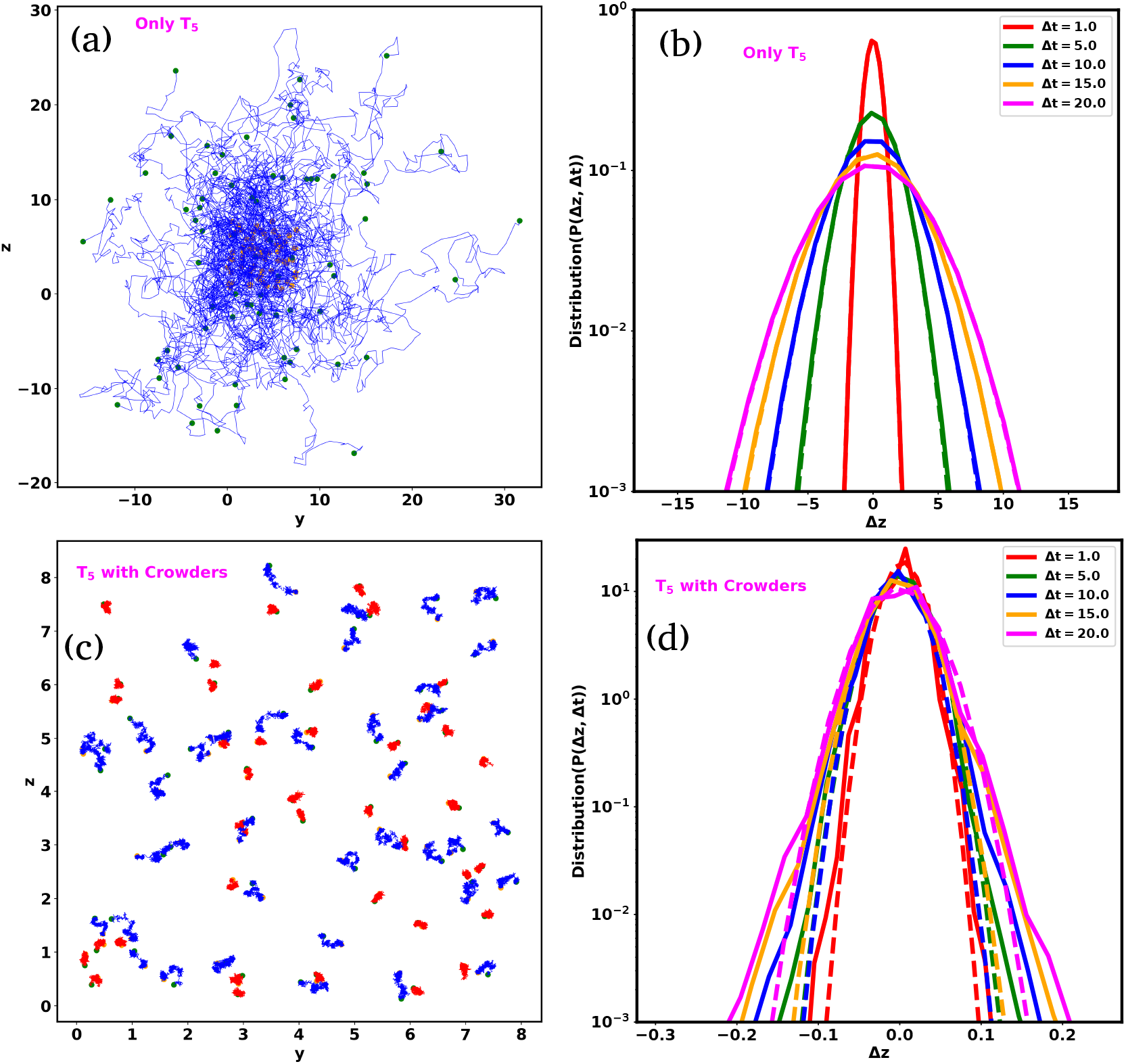
Comparison of Dynamic heterogeneity(DH) and non-Gaussian distribution between *T*_5_ particle. (a) The trajectory of (*T*_5_) particles in the absence of crowders. Each trajectory spans the same time period (6.0*τ*_*B*_). These particles move more quickly and uniformly, filling up more of the cytoplasm. (b) Distribution of displacement (Δ*z*) for *T*_5_ with different delay times Δ*t* in the absence of crowders. (c) The trajectory of (*T*_5_) in the presence of crowders. Each trajectory of *T*_5_ span the time period (1000.0*τ*_*B*_). However, from the variation of trajectory sizes of *T*_5_ within the same time period, we can argue that there are co-existence of fast (blue) and slow (red) moving particles, supporting the strong indication of dynamic heterogeneity. (d) Distribution of displacement (Δ*z*) for *T*_5_ with different delay times Δ*t* in the presence of crowders. In all figures (a and c), the start and end points of the respective trajectories are denoted by magenta and green colored scatter points. In all plots of (b) and (d), the simulated data is represented by solid lines, while the fits of a Gaussian function are depicted by dotted lines.

To quantitatively compare the observed dynamics with the expected dynamics of diffusive particles, we have examined the particle displacement distributions. Here, we have calculated the displacement of the *T*_5_ particles (z component) as 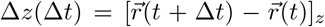, where 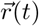 is the position of the particle at time *t* and Δ*t* is the delay time.

Figures 4(b) and Figures 4(d) illustrate the distribution of Δ*z* for *T*_5_ particles in the absence and in the presence of crowders respectively, with different delay time Δ*t*. In all these plots, the solid lines represent the simulated data, while the dotted lines depict the fits to a Gaussian function represented as 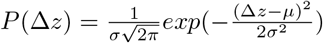, where *σ* and *μ* correspond to the standard deviation and mean of Δ*z*, respectively. When the tracer particles are placed in isolation, the distribution of displacment closely matches Gausssian distribution, as expected from Brownian motion. In the presence of the crowder the distributions deviate from the Gaussian function, which is another salient feature of glassy dynamics.

#### Box 1

Proteins translation requires the largest portion of metabolic energy. The protein translation machinery ribosomes are the most abundant proteins in bacterial cytoplasm. Ribosomes are large and undergo large change (10x) in diffusivity during the switch of their state from subunits to actively transcribing functional units in a polysome.

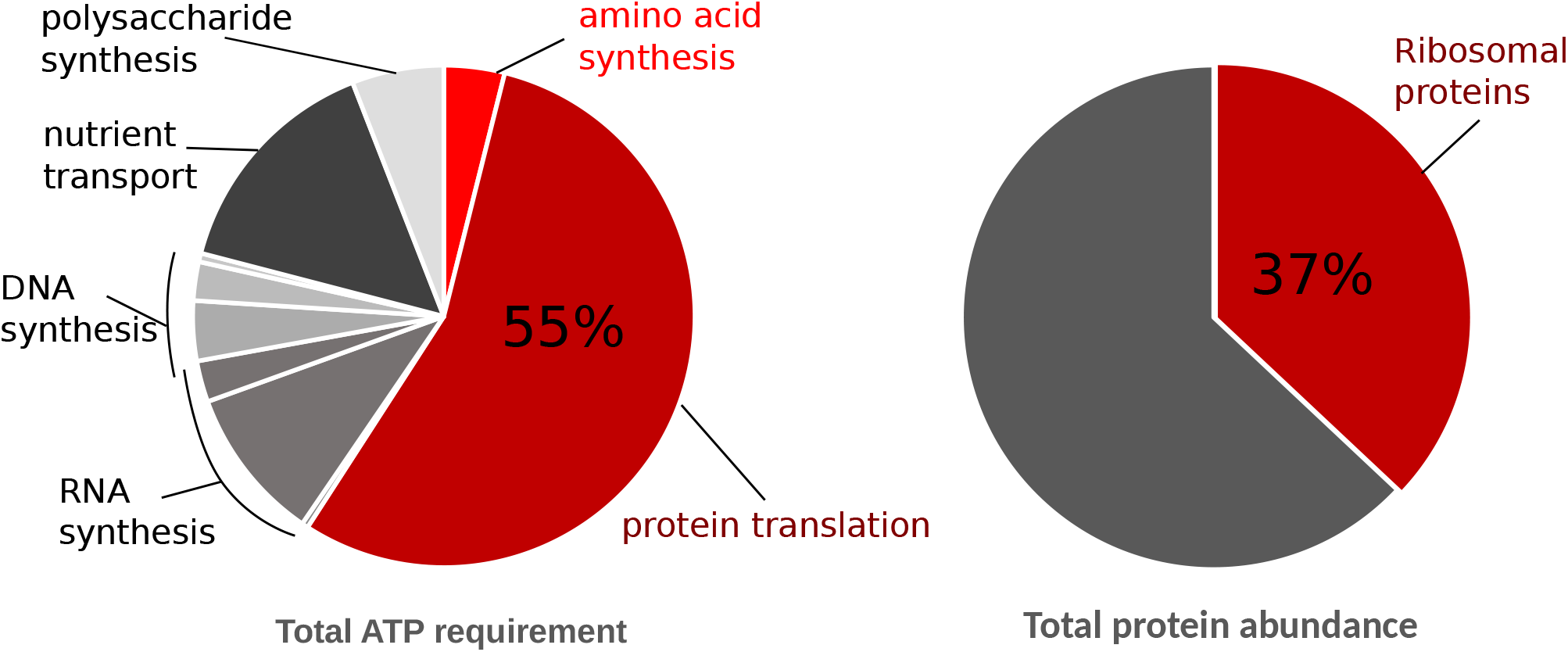

### E. Translation can fluidize the glassy cytoplasm

The experimental investigations by Weber et al. [13] and Parry et al. [12] have indicated that the metabolic activities of living cells can fluidize the glassy dynamics of large cytoplasmic particles. Specifically, the research from both groups has shown that the presence and abundance of ATP is a key determinant of this metabolic activity-dependent fluidization.

In living cells, a substantial portion of ATP is devoted to translation, specifically for aminoacylating tRNAs and regenerating GTPs. This energy expenditure is substantial, as translation activities consume approximately two-thirds of the total ATP pool within cells, significantly exceeding the demands of other energy-dependent cellular processes like nutrient transport, RNA, and DNA synthesis [19–22] (see Box 1). Consequently, translation emerges as a prime candidate whose rate and frequency could be profoundly influenced by ATP availability.

Interestingly, ribosomes, the essential machinery for translation, occupy a substantial portion of the cellular proteome and significantly contribute to cellular crowding. This crowding, in turn, governs the compaction of the nucleoid and the segregation of the remaining cytoplasmic components [8]. Any ATP-dependent alterations in ribosomal activity could lead to substantial changes in cytoplasmic crowding dynamics, potentially impacting the cytoplasm’s glassy nature.

What adds to the intrigue is the unique connection between ribosome mobility and their activity. Actively translating ribosomes form polysomes, groups of multiple ribosomes operating on the same mRNA strand. Polysomes exhibit slow diffusion within the cytoplasm and maintain strong segregation from the nucleoid. Conversely, freely diffusing ribosome subunits move rapidly and can traverse the nucleoid with ease. Notably, the diffusion rate of ribosome subunits is at least an order of magnitude faster than their actively engaged counterparts. The transition between these two states can lead to substantial alterations in the cytoplasm’s caging dynamics in an ATP-dependent manner. Taken together, these factors position the combination of ribosomes and translation activity as an exceptionally suited candidate for mediating metabolic activity-dependent fluidization of the cytoplasm.

We aimed to explore the impact of metabolic activity, particularly polysome-ribosome inter-conversion, on the diffusion of various biomacromolecules within *E. Coli* [23–25]. As illustrated in Figure 5(a), we modeled this inter-conversion as follows: at specific intervals *t*_*i*_, we randomly selected approximately *∼* 26% of polysomes for conversion, or ‘switching,’ into monomeric ribosomes. In our model, this state transition involved the removal and addition of a single 70S subunit from and to the polysome simultaneously. Figure 5(a) visually represents these switching schemes, where a 70S monomer dissociates into two subunits (30S and 50S) at one end of the polysome, while two subunits (30S and 50S) from the nearby cytoplasmic region join to the other end of the polysome to form a single subunit (70S). This process repeats after the same interval (*t*_*i*_). The corresponding switching rate is denoted as *k*_*s*_ = 1*/t*_*i*_ in units of 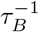 .

**FIG. 5.**
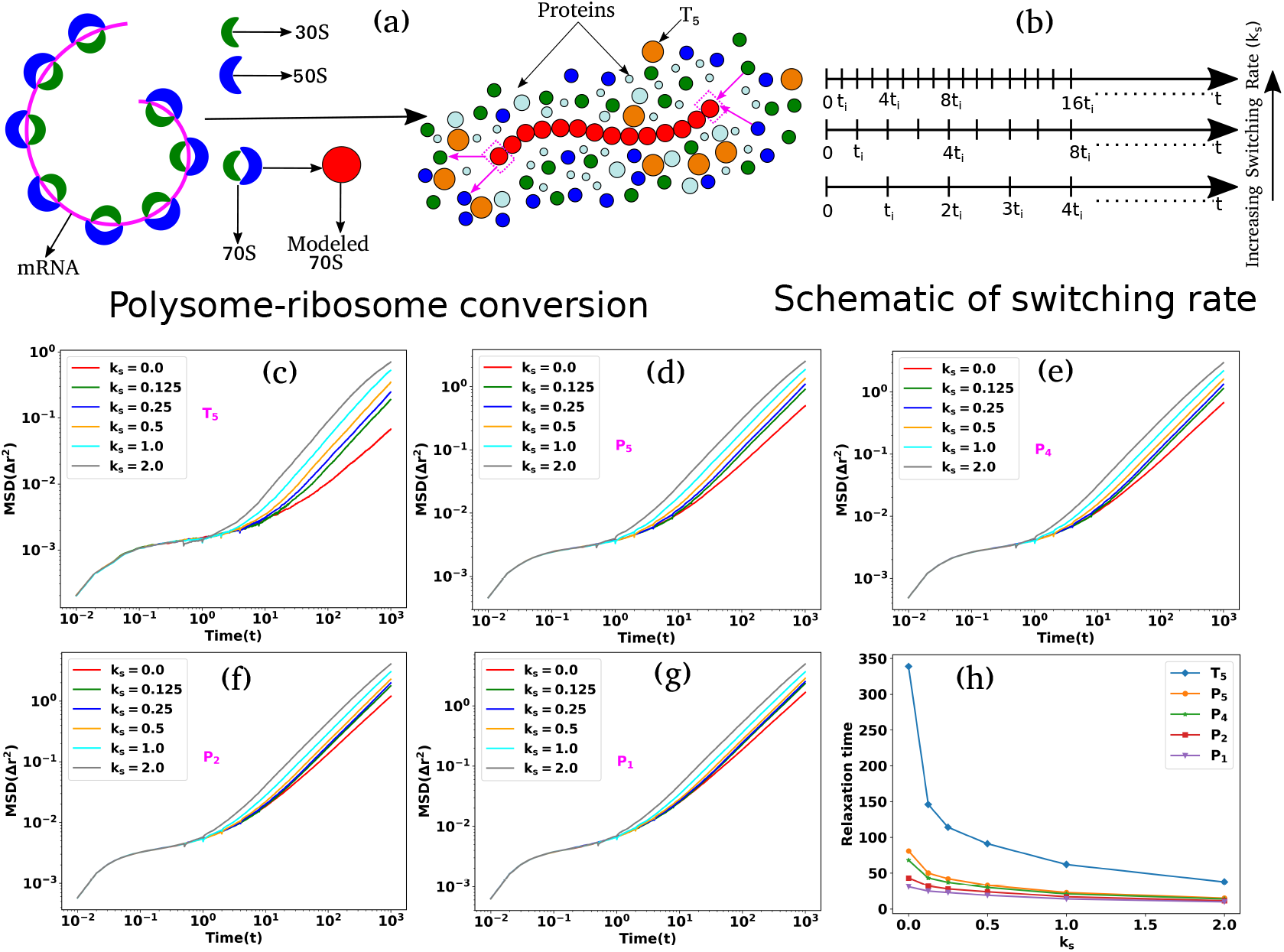
Translation activities can escalate the dynamics of cytoplasmic particles. (a) Schematic representation of polysome-ribosome conversion. The magenta color thread represents the mRNA chain, with the ribosomal subunits located along its length. Two ribosomal sub-units (30S and 50S) associate to create another sub-unit called 70S and polysome is the threads of 70S. The polysome-ribosome conversion has been modeled as follows: a 70S bead from one end of the polysome dissociates into two beads (30S and 50S) and two beads (30S and 50S) from the nearby region join to the other end by forming a single bead (70S). (b) Schematic representation of switching rate. The entire trajectory is represented by the time (0 *− t*), and in between switching happens at regular intervals of (*t*_*i*_). The switching rate will increase if there are more switching occurrences within the same interval. MSD as a function of time with various switching rates for different sizes particles in ascending order (c) (*T*_5_), (d) (*P*_5_), (e) (*P*_4_), (f) (*P*_2_) and (g) (*P*_1_) respectively. For all the particles, the MSD curves are showing ballistic, diffusive, and caging regions which are the hallmarks of the glassy dynamics and the length of cages diminishes with the increase in switching rate. In contrast, for the smaller particles(*P*_1_) and (*P*_2_), the switching effects are less prominent. (h) Structural relaxation time (*τ*_*α*_) as a function of switching rate (*k*_*s*_) for particles of different sizes. For the largest and intermediate-sized particles as the switching rate increases the relaxation time decreases. However, for smaller sizes particles the relaxation times are almost similar irrespective of the polysome ribosome switching.

Figure 5(b) presents a schematic representation of switching rates. The total trajectories cover the time span (0*−t*), with switching events occurring at regular intervals of (*t*_*i*_). Depending on various cellular factors such as ATP-mediated energy, ribosomal interactions, translation elongation velocity, and more [4, 26–29], the switching rate can vary. An “increasing switching rate” implies a higher frequency of switching events within the same time interval. The dotted lines suggest that there are additional switching events in the remaining intervals, which are not depicted in the diagram.

Figures 5(c-g) depict the Mean Squared Displacement (MSD) curves for various species as a function of time. These curves are generated for different switching rates (*k*_*s*_) for *T*_5_ and various protein sizes, arranged in ascending order of their diameter. In the absence of any switching, the MSD curve for all particle sizes exhibits distinct regions: a ballistic phase at shorter timescales, followed by a diffusive phase at longer timescales, with a plateau separating the two. This plateau signifies that the tagged particles are confined by neighboring particles, a hallmark of glassy dynamics.

Interestingly, the introduction of polysome-ribosome inter-conversion leads to a reduction in the duration of the plateau as the switching rate increases. Simultaneously, the diffusion of these particles accelerates, indicating a gradual improvement in cytoplasmic fluidity. However, these effects are less pronounced for smaller proteins (*P*_1_ and *P*_2_) compared to larger particles.

To gain deeper insights into the relationship between relaxation time and switching rate for particles of different sizes, we calculated the structural relaxation time (*τ*_*α*_) and plotted it as a function of *k*_*s*_ for different sizes of particles, as shown in Figure 5(h). The results revealed that as the switching rate increases, the relaxation time decreases for the largest and intermediate-sized particles. However, for smaller particles, the relaxation times remain almost the same, regardless of the switching rate. This finding indicates that the conversion between polysome and ribosome has more impact on the dynamics of larger particles, which display glassy dynamics in the absence of switching.

### F. High copy number and large molecular mass of ribosomes makes them effective disruptors

In the preceding section, we investigated the effects of switching dynamics on tracer particles with varying sizes. Now, we turn our attention to a different aspect: How does the size of the switching particle itself influence the degree of fluidization? To explore this question, we conducted a new series of simulations, manipulating the size of ribosomal sub-units (30S and 50S) and polysome particles (70S). The modular structure of our computational model affords us the flexibility to design such ‘control’ scenarios, allowing us to tune the size of the switching ribosomes.

Essentially, we designed two types of control simulations. In the first setup, we altered the size of the ribosomal sub-units and polysome particles to smaller dimensions (i.e., *σ*_30*S*_ = 12 nm, *σ*_50*S*_ = 15 nm, and *σ*_70*S*_ = 18 nm). In the second setup, we increased the size of the ribosomal sub-units and polysome particles (i.e., *σ*_30*S*_ = 19 nm, *σ*_50*S*_ = 22 nm, and *σ*_70*S*_ = 25 nm). To maintain a consistent packing fraction in the cytoplasm, as per the original simulation settings, we added or removed additional protein particles (*P*_1_-*P*_5_). These systems are denoted as ‘12-15-18’ and ‘19-22-25’ based on the sizes of the 30S, 50S, and 70S particles. The original system is referred to as ‘14-17-20’ according to the sizes used in the original system.

Figures 6(a) represent the MSDs of a the large tracer particle *T*_5_ as a function of time, for different combinations of switching particles. For each combination, we have compared the MSD trends at a switching rate 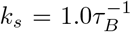 and with a switching rate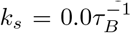 (no switching). We note that increasing the size of ribosome-polysome particles results in a notable change in dynamics. Specifically, as the size of the ribosome-polysome particles increases, the tagged particles (*T*_5_) show enhanced diffusion. Remarkably, in the case of the larger ribosome-polysome system (‘19-22-25’), the extent of the caging effect becomes less pronounced, leading to even higher diffusion compared to all other scenarios. In our simulation setup, the nonbonded interactions between all the particles are repulsive. Consequently, larger-sized particles experience stronger repulsive forces compared to their smaller counterparts. This result signifies the likely importance of the large size of ribosome particles in providing sufficient disruption to their local surroundings to cause fluidization.

**FIG. 6.**
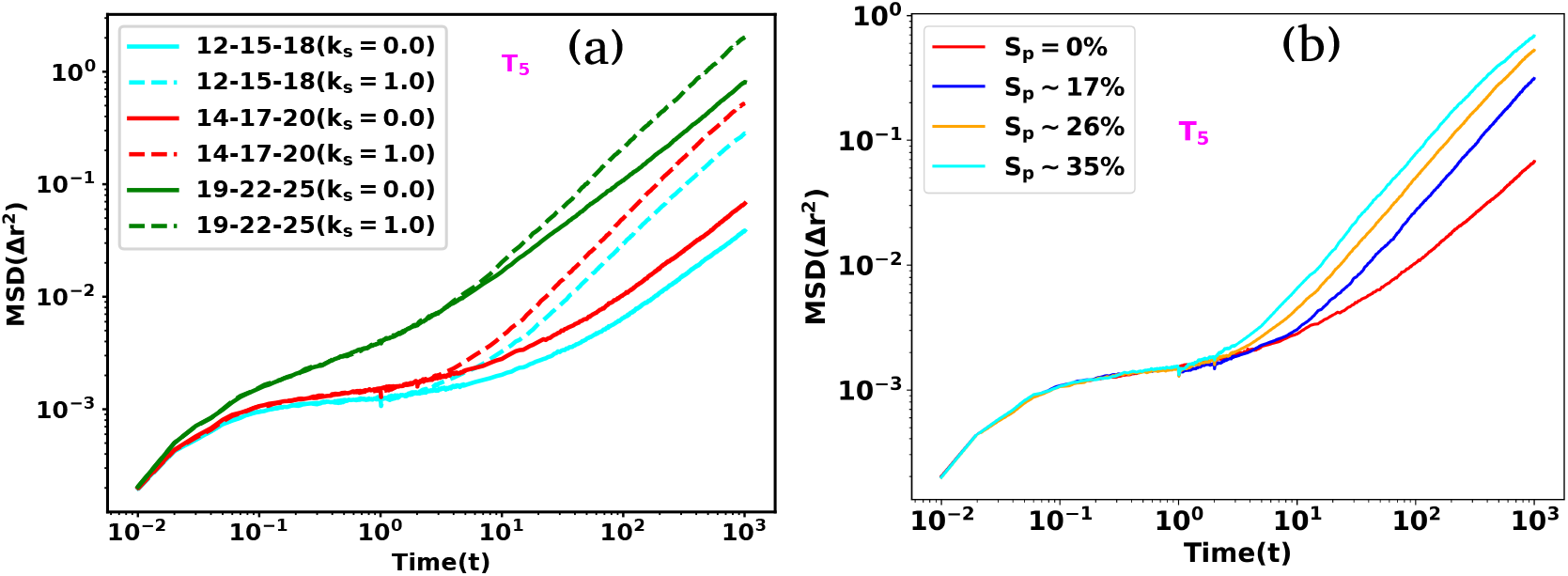
The impact of switching particle sizes. (a) MSD as a function of time is presented for polysome-ribosome systems of varying sizes, both with and without the presence of the switching effect. Notably, as the sizes of the switching particles increase, the tracer particles *T*_5_ exhibit enhanced diffusion. (b) MSD as a function of time for different percentages of polysome particle switching. As the percentage of switching (*S*_*p*_) increases the tracer particles *T*_5_ show a systematic escalation in their mobility.

Finally, we delved into the role played by the density of the disruptor moiety, exemplified by the ribosome, in the disruption of glassy states. To analyze this, we manipulated the fraction (*S*_*p*_) of polysomes undergoing state transitions within a fixed switching rate (*k*_*s*_ = 1.0) and observed its impact on the Mean Squared Displacement (MSD) of the tracer particle *T*_5_ (Figure 6(b). As anticipated, an increase in the fraction of switching particles led to a rapid escalation in the mobility of the tracer particle, concurrently diminishing the plateau associated with caging. In a metabolically active cell, a substantial number of ribosomes continuously transition between states, commencing or concluding rounds of translation. As one of the largest copy-number molecules housing an extensive molecular machinery within the cell, ribosomes disturb the local organization of crowders, facilitating mobility for other cellular machineries.

While the bacterial cytoplasm contains proteins with higher copy numbers, such as EFTu [30, 31], it is conceivable that EFTu molecules also undergo state transitions dependent on protein synthesis. This arises from their binding and unbinding interactions with translating ribosomes, potentially amplifying the effect of fluidizing the cytoplasm. The fraction or frequency of switching among translating ribosomes and their associated machinery is anticipated to be contingent on the cell’s metabolism, thereby presenting a mechanism for activity-dependent fluidization.

## CONCLUDING REMARKS

Our computational investigation of the bacterial cytoplasm model has provided critical insights into the origins and nature of the glassy behaviour of biomolecules within bacterial cytoplasm. Specifically, our model elucidates how glass dynamics can emerge in the apparently moderately crowded bacterial cytoplasm and why it is size-dependent, as has been experimentally observed in prior studies. In this paper, we find that the spatial organisation of bacterial cytoplasm is a critical underlying factor. This organisation creates regions of high crowding and exhibits a size-dependent arrangement of cytoplasmic particles, a phenomenon stemming from the sieve-like structure of the bacterial nucleoid.

Consistent with previous experimental studies, our model shows that the nucleoid is compacted due to the presence of crowding agents, such as a large number of ribosome particles [8–10]. As a consequence of this compaction, the ribosomes and similar large crowding agents are concentrated in the end-cap region of the cytoplasm. Consequently, the packing density in these areas is significantly higher than the average packing density of the cytoplasm as a whole. This localised increase in packing density leads to extreme crowding, which is the fundamental cause of the caging and resulting glassy dynamics observed in bacterial cytoplasm. Moreover, we demonstrate that the degree of segregation from the nucleoid and the enrichment in ribosome-rich regions depend on the size of the particles involved. As a result, the effect of crowding is more pronounced in the case of larger proteins, leading to size-dependent glassy dynamics, where larger proteins exhibit more significant effects.

Continuing our investigation, we next delve into the factors responsible for the metabolism-dependent fluidization of this glassy behaviour, a crucial mechanism for sustaining cell viability. We investigated the role of protein translation, which stands as the most energy-consuming process within the cell, in generating the necessary motion within the densely packed cytoplasm to liberate the confined particles. Specifically, ribosomes and the multitude of proteins associated with them undergo substantial changes in their dynamics when transitioning from an active translation state to a state of free diffusion. This frequent shift in the dynamic behaviour of the translation-related proteins, which represent a significant portion of the bacterial proteome, disrupts their local surroundings sufficiently to loosen the constraints on the caged particles, ultimately promoting fluidity in their immediate vicinity. Put together, our model provides evidence that protein translation could be the key energy-dependent process underlying the metabolic activity dependent fluidization of bacterial cytoplasm.

In essence, our research introduces a compelling concept: the spatial organisation of the bacterial cytoplasm, which forms the foundation of glassy dynamics, is primarily orchestrated by the entropic segregation of ribosomes from the bacterial chromosomal DNA. This segregation likely plays a critical role in localising activities to increase concentration and reduce search times [32]. Intriguingly, the activity of the very same ribosomes serves as a dynamic force that disrupts the system, alleviating the confining effects. This activity-driven fluidization is of paramount importance for preserving the enzymatic functions of the cell. It’s noteworthy that at the heart of this crucial fluidization process lies one of the most ancient enzymatic mechanisms in evolution: protein translation.

## MATERIALS AND METHODS

### Model and Method

The cytoplasm of *E. Coli* is a highly complex environment and the exact constituents are still unknown. However, it consists of DNA, RNA, polysomes, ribosomes, and numerous poly-disperse proteins, etc. Here we have modeled the *E. coli* chromosome as a bead-spring polymer chain with each bead representing 5 *×* 10^3^*basepair* (5 kbp) and encodes Hi-C-derived contact probability matrix [15], as described in our previous works [33–36]. In brief, the bonded interactions between adjacent beads have been modeled by harmonic springs with a spring constant *k*_*spring*_ = 300*k*_*B*_*T/σ*^2^, and the non-bonded interactions are modeled by the repulsive part of Lenard Jones (LJ) potential i.e., *V*_*nb*_(*r*) = 4*e*(*σ/r*)^12^ with diameter and mass given by *σ*, and *m* respectively (in real units *σ* = 67.31 nm and *m* = 3245*kDa*). The Hi-C-derived inter-genomic contacts are modeled as harmonic springs with probability-dependent distance and distance-dependent force constants (see SI method section). Unlike our previous model [33, 34], in our current study, we have incorporated polysome and ribosomes. Each 30S, 50S, and 70S ribosomal sub-units have been modeled as spherical particles with different diameters (*σ*_30*S*_ = 14 nm, *σ*_50*S*_ = 17 nm, and *σ*_70*S*_ = 20 nm) and each polysome is modeled by 13-mer of 70S ribosomal sub-units which behave like a freely jointed chain. To investigate the spatial arrangement of DNA and ribosomal subunits, we conducted simulations within a sphero-cylindrical confinement that emulates the cell wall. In this simulation environment, all cytoplasmic particles interact through excluded volume interactions, and their behavior is described by the repulsive component of the Lennard-Jones potential, with the exception of DNA and ribosomal subunits. Our observations indicate that a modest attractive interaction between DNA and ribosomal subunits is required to achieve a simulated linear density that closely approximates experimental data. This finding implies that cytoplasmic crowders exhibit some level of non-inert behavior. Detailed values for the parameters *e* and *σ* for all particles can be found in the Supplementary Information (SI).

In a similar way, we have also introduced 2000 copies of tracer particles (*T*_1_ to *T*_5_) one at a time in ascending order of their diameter (10 *−* 50) nm. These tracer particles engage in interactions among themselves as well as with the other cytoplasmic components, characterized by excluded volume interactions governed by the repulsive component of the Lennard-Jones potential. Similar to polysomes, large tracers also exhibit pronounced segregation from the nucleoid, with the extent of segregation dependent on their size.

The bacterial cytoplasm not only contains polysomes and ribosomes but there are also numerous other protein particles. To delve into the impact of molecular crowding on the length scale of the whole cell, the number of moieties will be huge. Thus, our focus is directed toward analyzing the dynamics of particles within the end-cap region, where a majority of polysomes and larger tracer particles reside. To initiate this study, we initially quantified the number of 30*S*, 50*S*, 70*S*, and *T*_5_ particles within the end cap regions from our previous simulations. These particles were subsequently placed within a cubical box, its volume comparable to that of the cell’s end cap. Additionally, to achieve a desired packing fraction (*pf* = 0.57), we introduced five distinct types of polydisperse protein particles (*P*_1_ *− P*_5_) with sizes ranging from (13 *−* 18) nm. These particles also interact with each other and the rest of the cytoplasmic particles via the repulsive component of the Lennard-Jones potential.

All the components of our model follow the Brownian dynamics and we performed the simulations in reduced units using open-source Software GROMACS 5.0.6 [37] with simulation time step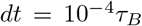. We have modified the source code to introduce sphero-cylindrical confinement. Here all simulation length, time, and energy are in the unit of *σ* (the diameter of the DNA bead), *τ*_*B*_ and *k*_*B*_*T* respectively, and we have kept the *k*_*B*_*T* = 0.55. The mass, diameter, and the details modeling aspect of each component are given in SI(Table-S1). We mapped the reduced unit of simulation to the physical units by taking the real value of the diameter of each DNA bead (*σ* = 67.31 nm) [34], temperature (*T* = 303*K*), and viscosity of the cytoplasm (*η* = 17.5*Pa*.*s*) [38, 39]. In real physical unit the value of *τ*_*B*_ becomes 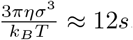 .

## DYNAMIC QUANTITIES

### Mean Squared Displacement (MSD)

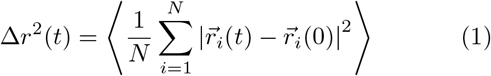

where 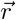, and N are the position of the particle and number of particles respectively and ⟨ .. ⟩ represents the ensemble average.

### Structural Relaxation Time (*τ*_*α*_)

Generally, relaxation time is calculated from a twopoint density-density auto-correlation function, defined as

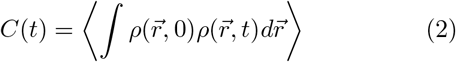

Where 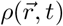 is the density at position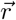 at time t and ⟨… ⟩ is the ensemble average. But, here we have used a popular variant of *C*(*t*), which is known as the overlap function and is defined as [40–42]

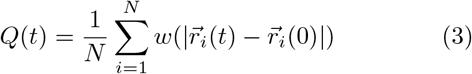

where *w* is the window function, *w*(*r*) = 1 if *r < a*_0_ and 0 otherwise, and *a*_0_ is the cutoff distance at which the root mean squared displacement (RMSD) of the particles exhibits a plateau. In our study, we have chosen the *a*_0_ = 0.115*σ*, to neglect small de-correlation because of the vibration of the particles in their respective cages. The structural relaxation time *τ*_*α*_ is defined as *Q*(*t* = *τ*_*α*_) = 1*/e*.

## ACKNOWLEDGMENTS

We thank Saroj Kumar Nandi and Smarajit Karmakar for useful discussions. We acknowledge the computation facilities provided by TIFR Centre for Interdisciplinary Sciences, India. We acknowledge the support of the Department of Atomic Energy, Government of India, under Project Identification No. RTI 4007 and Core Research grants provided by the Department of Science and Technology (DST) of India (CRG/2023/001426) to J.M. The work by S.B. was supported by the Wellcome Trust Award [grant number RG89305].

