## Supplemental Method and tables for "Protein Translation Can Fluidize Bacterial Cytoplasm"

### Supplementary Information
### (Protein Translation Can Fluidize Bacterial Cytoplasm)

Palash Bera,[1] Abdul Wasim,[1] Somenath Bakshi,[2,*] and Jagannath Mondal[1,†]

[1] *Tata Institute of Fundamental Research, Hyderabad 500046, India*

[2] *Department of Engineering, University of Cambridge, Cambridge, United Kingdom*

*

†

#### A. Model and Simulation Method Details

We model the DNA of *E. Coli* as a bead-in-a-spring polymer chain, with each bead representing $5 \times 10^3$ bp (5 kb) and this resolution is the same as the Hi-C interaction maps reported by Lioy et al. The diameter and mass of each bead are $\sigma$ and $m$ respectively. The adjacent beads are interacting harmonically with spring constant $k_{spring} = 300 k_B T/\sigma^2$. The non-bonded interactions of the beads have been modeled by the repulsive part of Lenard-Jones potential i.e $U_{nb} = 4\epsilon(\frac{\sigma}{r})^{12}$. The Hi-C contacts are also modeled by harmonic springs which act as cross-links between different beads of DNA. We converted the Hi-C probability matrix to the distance matrix as

$$D_{ij} = \sigma/P_{ij} \tag{1}$$

where $i$ and $j$ are the row and column index of the matrix respectively. The Hi-C restraining potential between a pair of Hi-C contacts at a separation of $r_{ij}$ is defined as

$$U_{Hi-C}(r_{ij}) = \frac{1}{2} k_{ij} (D_{ij} - r_{ij})^2 \tag{2}$$

where $k_{ij}$ is the distance-dependent force constant that can be calculated as

$$k_{ij} = k_0 e^{-\frac{(D_{ij}-\sigma)^2}{w^2}} \tag{3}$$

where $k_0$ is the upper bound of the spring constant and $w^2$ is a constant value. In our simulation we have kept the value of $k_0$ and $w^2$ same as of our previous study i.e $k_0 = 10 k_B T/\sigma^2$ and $w^2 = 0.3$.

Apart from DNA, bacterial cytoplasm consists of polysome, ribosomes, and numerous other poly-disperse protein particles. There are mainly two types of ribosomal sub-units, 50S(large) and 30S(small). During the translation, two sub-units of ribosomes attach together to form 70S ribosomes and the threads of 70S ribosomes are called a polysome. Generally in living cells, there are 80% of the ribosomes are in the form of polysome. We model the ribosomes (30S, 50S, and 70S) as spherical particles with different masses and diameters. The polysome has been modeled by 13-mer of 70S ribosomal sub-units. The adjacent beads of the polysome interact harmonically with spring constant $K_{polysome} = 17000 k_B T/\sigma^2$. Previous experimental studies showed that there are $\sim$ 26000 ribosomal sub-units and among them, 80% ribosomes are in the form of polysome. Thus in our simulation, we have incorporated the same number of spherical particles corresponding to 30S 50S, and 70S ribosomal sub-units. Among them 2600 particles are the 30S, 2600 particles are 50S and 20800 particles are the 70S. As we model polysome as 13-mer of 70S ribosomal sub-units, the total number of polysome chains is 1600. The polysomes, ribosomes, and protein particles are interacting with each other and with DNA repulsively and the repulsive interactions are taken from the repulsive part of LJ potential. Our observations indicate that a modest attractive interaction between DNA and ribosomal subunits is required to achieve a simulated linear density that closely approximates experimental data. For the nonbonded interactions between DNA and ribosomal subunits, we employed the complete Lennard-Jones (LJ) potential, denoted as $U_F$, defined as: $U_F = 4\big(\epsilon^r_{ij}(\frac{\sigma_{ij}}{r_{ij}})^{12} - \epsilon^a_{ij}(\frac{\sigma_{ij}}{r_{ij}})^6\big)$. Here, the subscripts $i$ and $j$ represent particle indices while $\epsilon^r_{ij}$ and $\epsilon^a_{ij}$ correspond to the repulsive and attractive components, respectively. Specifically, $\epsilon^r_{ij} = 1$ for all particles, and $\epsilon^a_{ij}$ assumes a value of 0.2 exclusively for interactions between DNA and ribosomal subunits. This parameterization signifies a relatively weak attractive interaction between these two types of particles. Apart from bonded and non-bonded interactions, all the particles are confined within a spherocylindrical of length $L = 45.754\sigma$ and diameter $d = 12.181\sigma$, mimicking the cell wall. The confinement potential is defined as

$$U_{res}(r, R_0) = \frac{1}{2} k_{res} \left|\vec{r} - \vec{R}_0\right|^2 \Theta \left|\vec{r} - \vec{R}_0\right| \quad (4)$$

| Particle Type | Number | mass in m<br>$m = 649 \times 5 = 3245$kDa | diameter in $\sigma$<br>$\sigma = 67.31nm$ |
|---|---|---|---|
| DNA beads | 928 | 1 | 1 |
| 30s | 257 | 0.26 | 0.20 |
| 50s | 267 | 0.42 | 0.25 |
| Polysome-beads (70s) | 173 × 13 | 0.71 | 0.29 |
| $P_1$ | 2308 | 0.24 | 0.19 |
| $P_2$ | 4886 | 0.27 | 0.21 |
| $P_3$ | 6274 | 0.29 | 0.23 |
| $P_4$ | 4886 | 0.31 | 0.25 |
| $P_5$ | 2308 | 0.34 | 0.27 |
| $T_5$ | 305 | 0.94 | 0.74 |

TABLE S1. Different types of particles with their number, mass, and diameter for simulations inside the cubical box. Here DNA beads serve as the reference particles, where all measurements of diameter and mass are expressed in terms of DNA particles.

where $R_0$ is the center of the spherocylinder and $k_{res}$ is the spring constant that controls the softness of the confinement. Here we have used $k_{res} = 310k_BT/\sigma^2$. The $\Theta$ is the step function that will activate if any particle gets out from the confinement. So the total configurational potential energy is given by

$$U_{tot} = U_b + U_{nb} + U_{Hi-C} + U_{res} + U_F \tag{5}$$

where $U_b$, $U_{nb}$, $U_{Hi-C}$, and $U_{res}$ are the bonded, non-bonded, Hi-C restraining, and confinement restraining potential.
